# Hotair and Malat1 Long Noncoding RNAs regulate Bdnf Expression and Oligodendrocyte Precursor Cells Differentiation

**DOI:** 10.1101/2021.08.09.455020

**Authors:** Fatemeh Khani-Habibabadi, Leila Zare, Mohammad Ali Sahraian, Mohammad Javan, Mehrdad Behmanesh

## Abstract

BDNF has remarkable protective roles in the central nervous system to ensure neurons and glial cells survival and proper functions. The regulatory processes behind the BDNF expression have not been revealed completely. Here, it was explored whether *Malat1* and *Hotair* lncRNAs play roles in the regulation of *Bdnf* expression level, modification of fingolimod downstream pathway, and oligodendrocytes precursor cells maturation. By *Hotair* and *Malat1* downregulation, their regulatory mechanism on *Bdnf* expression was investigated. Immunostaining and RT-qPCR assays were employed to assess the effects of fingolimod and lncRNAs on OPCs maturation. The results represented that *Hotair* and *Malat1* lncRNAs may regulate Bdnf expression in primary glial cells significantly, and also can coordinate fingolimod stimulatory effect on *Bdnf* expression. Furthermore, *Malat1* may have a role in the last stages of the intrinsic oligodendrocyte myelination. Here it was demonstrated that these lncRNAs have critical roles in the *Bdnf* level, fingolimod mechanism of action, and OPCs maturation. Understanding the regulatory mechanism of neurotrophins leads to a better comprehension of the neurodegenerative disorders pathogenesis and designing more effective treatments.

## 1. Introduction

Brain-derived neurotrophic factor (BDNF) is a member of the neurotrophins family with some well-known roles such as regulation of neural development, differentiation, survival, and synaptic plasticity. There are two types of secreted BDNF, pro-BDNF and mature-BDNF [1]. By binding to the P75 receptor, pro-BDNF stimulates pro-apoptotic effects, while mature-BDNF triggers pro-survival pathways through binding to the TrkB receptor [2]. By BDNF production, astrocytes, microglia, brain-resident macrophages, and infiltrated lymphocytes in the brain lesion can protect neural survival [3, 4]. It is well demonstrated that BDNF induces oligodendrocyte precursor cells (OPCs) regeneration, and differentiation [5, 6].

In the central nervous system (CNS), oligodendrocyte cells are responsible for myelin production. As immune cells cross the blood-brain barrier (BBB) and ignite an inflammatory environment, oligodendrocytes, which are the most sensitive glial cells to inflammatory conditions, undergo apoptosis that results in myelin loss and plaque formation [7]. Damaged myelin leads to lesions and plaque formation, which interfere with the regular signal transmission and prone axons to the inflammatory milieu [8]. It has been suggested that myelin repair has two general steps, OPCs recruitment to the lesion region and differentiation into mature oligodendrocytes. Defects in any of these two pathways have been reported in demyelinating disorders. The remyelination process can occur to some extent, but its rate and efficiency decrease over time [9].

A destructive-protective role has been considered for astrocytes in demyelinating disorders. It is suggested that the inflammatory conditions in the CNS affect the astrocytes activation and genes expression profile. Severe astrogliosis prevents CNS healing by releasing more inflammatory cytokines; however, mild astrogliosis triggers the production of protective and growth factors such as BDNF, which protects newly synthesized myelin proteins [10, 11]. Like astrocytes, growing evidence shows the heterogeneous nature of microglia in this phenomenon. By activation, microglia can be developed into two distinct phenotypes, M1 and M2 in which, M1 microglia play a cytotoxic role by producing inflammatory factors such as IL-6, IL-1β, and TNFα in the advanced stages of the disease. However, the M2 phenotype is more prominent in the early stage and plays roles by producing protective factors such as BDNF and IGF-1 [12].

Fingolimod acts as a S1P agonist and targets the signaling pathway of the sphingosine-1-phosphate GPCR receptor. Phosphorylated fingolimod can bind to all types of S1PR except S1P2 and cause ubiquitination, internalization, and degradation of the receptor. Hence, lymphocytes do not respond to the S1P ligand gradient and get trapped in the lymph nodes [13]. It is well declared that through binding to S1P receptors on microglia, oligodendrocytes, astrocytes, and neurons, fingolimod plays protective roles in the CNS, including cell survival, OPCs differentiation, myelin repair, restoring synaptic function, a reduction in the secretion of inflammatory cytokines, and an increase in the neurotrophins production [14, 15].

Long non-coding RNAs (lncRNAs) are a class of non-coding RNAs that are longer than 200 nucleotides and cannot encode proteins [16]. Current studies have shown the important roles of lncRNAs in different human diseases such as neurodegenerative and demyelinating disorders [17]. In our previous study by bioinformatics analysis, it was demonstrated that there is a positive correlation between *BDNF* gene expression and two lncRNAs *HOTAIR* (HOX Transcript Antisense RNA) and *MALAT1* (Metastasis Associated Lung Adenocarcinoma Transcript 1) in neurodegenerative disorders [18].

It has been reported that *HOTAIR* lncRNA binds to the PRC2 and LSD1 complexes and directs them to their target loci resulting in gene silencing. Besides, *HOTAIR* can sponge miRNAs and inhibit their binding to their downstream target genes [19]. *MALAT1* lncRNA, also known as *NEAT2*, is one of the most abundant nuclear lncRNAs and is highly conserved among species. Like *HOTAIR*, *MALAT1* can regulate target genes through two distinct mechanisms: sponging their regulating miRNAs and conductance of the PRC2 complex, which induces histones methylation and gene suppression [20].

LncRNAs research has been becoming a hotspot in neuroscience; however, a few studies have considered their roles in the glial cells function and OPCs maturation. This study employed a combination of in silico, molecular and cellular techniques, including homology analysis, gene downregulation, primary glial culture preparation, immunocytochemistry, and RT-qPCR, to decipher the potential regulative roles of *Hotair* and *Malat1* lncRNAs in the *Bdnf* expression in glial cells and oligodendrocytes myelination. The obtained results demonstrate that Hotair and *Malalt1* lncRNAs may significantly regulate *Bdnf* expression in glial cells. Also, *Malat1* can plays a role in the myelination process. These data pave the way to a better illustration of networks behind the OPCs differentiation and can serve as possible targets for future therapies of demyelinating disorders.

## 2. Materials and methods

### 2.1. Preparation of mixed glia culture

Primary glial cells were collected from P_0_ Wistar rat pups. Briefly, six rat pups’ brain was dissected mechanically by scalpel blade and passed through 40 μm cell strainer (SPL Life Sciences, Korea). The cell suspension was then cultured on Poly-L-lysine (PLL)-coated (Sigma, USA) T75 cell culture flasks in DMEM High Glucose, FBS 10%, supplemented with 50 U ml^-1^ penicillin and 50 μg ml^-1^ streptomycin (Gibco) at 37 °C under a humidified atmosphere containing 5% CO2. Every 2-3 days the medium was changed completely until the culture reached to ideal confluency [21]. The procedures of this research were approved by the ethics community of Tarbiat Modares University (ID: IR.TMU.REC.1396.607).

### 2.2. OPCs isolation and differentiation

Approximately after 10 days, the primary glial culture reach to enough confluency with small sparkling OPCs on the top of the astrocytes layer. In this study, the OPCs were isolated for further analyses of fingolimod effects on oligodendrocytes maturation by the shaking method. This isolation method was based on differential adherent properties of glial cells. For this purpose, the flasks were shaken for 1 hour at 200 rpm to detach and discard microglia cells. Then, to detach OPCs from the astrocyte layer the flasks were incubated at 37 °C and 5% CO_2_, while shaking at 220 rpm overnight. The cell suspension was transferred by a pipette to an untreated petri dish and incubated for 30 minutes to gain better OPCs isolation by differential adhesion properties of contaminating astrocytes and microglia [21]. Unattached OPCs were centrifuged at 1200 rpm for 5 min. Then, 10,000 OPCs were seeded in 4-well PLL-coated plates in proliferative medium, containing DMEM/F-12 medium, L-Glutamax (Gibco, USA) and Sodium pyruvate (Sigma, USA), supplemented with 20 ng/mL human recombinant basic fibroblast growth factor (bFGF; Invitrogen, USA), 20 ng/mL epidermal growth factor (EGF; Invitrogen, USA), and 20 ng/mL Platelet-derived growth factor (PDGF, Invitrogen, USA) for 3 days at 37 °C under a humidified atmosphere containing 5% CO2. To induce OPCs differentiation, the proliferation medium was replaced with the differentiation medium including B-27 (Gibco, USA) supplemented with IGF-1 (10 ng/ml, Invitrogen, USA) and T3 (40 ng/ml, Invitrogen, USA) instead of growth factors, and till the end of the differentiation process, half of the differentiation medium was refreshed every other day [21, 22].

### 2.3. Fingolimod treatment

To explore the fingolimod (Sigma, USA) cytotoxicity effects, the glia cells were treated with different concentrations of fingolimod (0 nM, 5 nM, 25, 50 nM, 75 nM, and 100 nM diluted in HPLC-grade ethanol). Based on obtained result of cytotoxicity assay concentration of 20 nM fingolimod was chosen for treating of Primary glial culture and OPCs gene expression profile, proliferation, and differentiation.

### 2.4. Immunostaining

For immunostaining, the medium was removed, and the cells were fixed by paraformaldehyde for 20 minutes incubation at room temperature. To increase cell permeability, Triton 0.2% and 0.5% was used for cytoplasmic markers (GFAP, Iba1, PAX6, and PLP) and nuclear markers (Olig2), respectively. Then, the first antibodies, including Anti-MBP antibody (1:200, Santa Cruz sc-271524, USA), Anti-Olig2 antibody (1:200, Abcam Inc. ab9610, USA), Anti-GFAP antibody (1:300, Dako Z0334, USA), and Anti-Aba1 antibody (1:200, Wako 019-19741, USA) were dissolved in NGS 10% and incubated overnight at 4 °C. The Alexa Fluor^®^ 568 secondary antibodies, including goat anti-mouse, and goat anti-rabbit antibodies (1:1000, Life Technologies, A11004 and A11036, USA), were used, and Nuclei were stained by 0.5 /g / ml DAPI [23]. The fluorescent signals were detected by a fluorescent microscope (Olympus, Japan).

### 2.5. Cell Proliferation and Viability Assay

The AlamarBlue^®^ assay was employed to measure fingolimod effects on cells proliferation, survival, and cytotoxicity. For this purpose, OPCs were treated with different concentrations of drug fingolimod. After 4 hours, the adsorption was measured at 540 and 630 by an ELISA reader (BioTek, USA) [24].

### 2.6. Gene expression analyses

Cell total RNA was extracted using the RiboEX solution (GeneAll, Seoul, Korea), and its quality and quantity were measured by spectrophotometry and 1.5% agarose gel electrophoresis. DNase I (Fermentas, USA) treatment was performed on RNA samples to remove any genomic DNA contamination. 3 μg of total RNAs was applied to synthesize the first strand of cDNA using M-MulV reverse transcriptase (Thermo Scientific, USA) by oligo dT, and random hexamer as primers (MWG, Germany) according to the manufacturer’s instruction. Gene expression analysis was performed using a StepOne™ PCR system (Applied Biosystem/MDS SCIEX, Foster City, CA, USA), with 10 ng cDNA, 5 μl of 2X SYBR^®^ Green master mix (Solis BioDyne, Estonia), and 4 pM of each forward and reverse primers up to final reaction volumes of 10 μl. Differential gene expression analysis was assessed using the 2^-ΔCT^ method [25]. The relative expression of *Bdnf* (forward: 5-AGGGAGTGAAGATACCATCAGC-3 and reverse: 5-TATGAGAGCCAGCCACTGACC-3), *Hotair* (forward: 5-GATTTCAAACTGAGCCCCGATG-3, reverse: 5-CAGGTTTAAGAGAGCCACCCAA-3), and *Malat1* (forward: 5-ACAAGACCACAGCTCAGTAGC-3, reverse: 5-CCAAGTCTGCTATGTCCACCTG-3) genes were normalized to the housekeeping gene (actin, beta) *Actb* (forward: 5-ATGTGGATCAGCAAGCAGGA-3, reverse: 5-AAAGGGTGTAAAACGCAGCTC-3) (Metabion, Germany). With the help of homology search in sequence databases of mice and rats’ genome, the sequences of target genes were found and specific primers based on them were designed. All of the samples were tested at least in duplicate and the specificity of qPCR reactions was verified by melting curve analysis and running of amplification products on 12% polyacrylamide gel electrophoresis.

### 2.7. Gene silencing with DNAzyme

In this study, a single-stranded DNA sequence with an approximate length of 35 nucleotides, called DNAzyme, was used to reduce the expression of target genes [26]. Oligo 7 (DBA Oligo, USA), SnapGene (GSL Biotech LLC, USA) software, and NCBI-Blast were used to design these oligonucleotides based on the homology searched sequenced. Also, secondary structures formation within the target region of RNAs was checked by the RNAfold webserver [27]. To down-regulate lncRNAs, The *Hotair* DNAzymes (5-GGCATTGGTAAGGCTAGCTACAACGAGACACTCTCT-3 and 5-GAAATAGCCAGGCTAGCTACAACGATGTGTCTCT-3) and *Malat1* DNAzymes (5-CCTCATTCCCAGGCTAGCTACAACGACCAGCATTTG-3 and 5-GCAATCCCAGGCTAGCTACAACGACCCAACAGCTT-3) were used (Metabion Company, Germany). Then, the cells were transfected with 5nM DNAzymes by INTERFERin^®^ reagent (Polyplus, USA) according to the manufacturer’s instructions. After 24 and 48 hours of incubation, the total RNA and protein were extracted and kept in −80 ° C till further analyses. The whole of experiment was repeated at least twice.

### 2.8. Western Blotting

Cells were lysed in RIPA buffer (Cyto Matin Gene Co., Iran) according to the manufacturer’s instructions. 20 μg of total cell proteins were run in a 12% SDS-polyacrylamide gel and blotted to the PVDF membrane (Bio-Rad, USA). Membranes were incubated overnight at 4 °C with mature BDNF primary antibody (ThermoFisher OSB00017W, USA) followed by an hour incubation with HRP goat anti-rabbit secondary antibodies (Santa sc-2004, USA) at room temperature. After chemiluminescence visualization, reprobing procedure was done by 0.2 M NaOH for 5 minutes. Next, beta-actin antibody (Santa Cruz sc-47778, USA) as the internal control was applied on the membrane at 4 °C overnight, followed by an hour incubation with HRP goat anti-mouse secondary antibody (Santa Cruz sc-2005, USA) at room temperature [28]. Blots were visualized by chemiluminescence (ECL; Cyto Matin Gene, Iran) and the obtained bands were quantified using ImageJ software.

### 2.9. Statistical analyses

For two-group and multiple-group analyses, comparisons were applied by unpaired t-test and ordinary one-way ANOVA with Tukey’s multiple comparisons test, respectively. For cell survival assay, ordinary two-way ANOVA was employed with Dunnett’s multiple comparison test. All of the statistical analyses and graphs visualizations were accomplished by Prism – GraphPad 7 (GraphPad Software, USA). Error bars represent standard deviation, and the significance level was assigned as p < 0.05 (two-tailed test).

## 3. Results

### 3.1. Fingolimod treatment did not affect glial cell survival

To analyze the fingolimod effects on the glial cells, the primary glial culture was obtained from the rat pups’ brain (Fig. 1, a). The immunostaining results demonstrated the presence of Iba1^+^ cells (Microglia, Fig. 1, b), GFAP^+^ cells (Astrocytes, Fig. 1, c), Olig2^+^ (Oligodendrocytes lineage, Fig. 1, d), and MBP^+^ cells (mature oligodendrocytes, Fig. 1, e). Signal quantification for each specific cell marker antibodies determined that the microglia are the most abundant cell type in the culture (54%). OPCs and astrocytes consist about 27% and 17.5% of cells in the primary culture, respectively; however, the mature oligodendrocytes were scarce (0.85%). The presented results are obtained from at least three times independent biological repeated (Fig. 1, f).

**Fig. 1.**
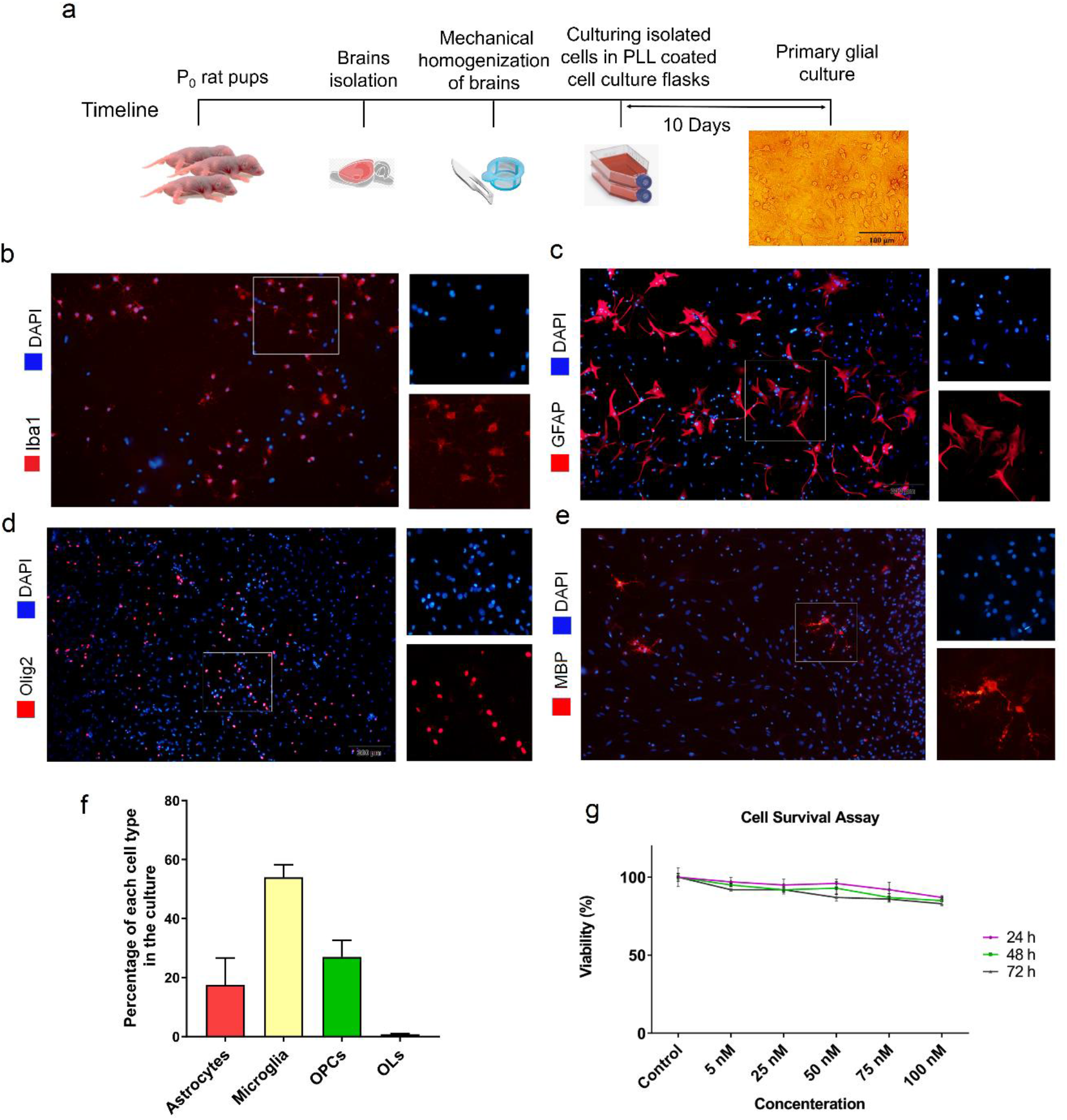
Preparation of primary glial culture from P_0_ rat pups’ brain. a, represents the timeline that was followed to make the primary glial culture. The immunostaining demonstrates the presence of b, Iba1^+^ marker of microglia; c, GFAP^+^ marker of astrocytes; d, Olig2^+^ marker of oligodendrocyte lineage; and e, MBP^+^ maker of mature oligodendrocyte cells in the prepared primary culture (scale bar= 200 μm, n=3). f, The quantification of glial cell lineage in the primary culture showed that oligodendrocyte lineage made up about 27% of the total population (error bar= standard deviation). g, Cell survival assay demonstrated that different concentrations of fingolimod treatment on the glial cells have no toxic effects at low doses; however, the cell viability reduced significantly by increasing the medicine concentration (*p< 0.05, **p < 0.01, ***p < 0.001, ****p < 0.0001, one-way ANOVA with Tukey’s multiple comparison test, error bar= standard deviation, n = 3). Abbreviation: PLL= Poly L-lysin. The experiment repeated three times, and the significance level was assigned to 0.05.

Proliferation assay demonstrated that fingolimod treatment had no cytotoxic effects on glial cells at 5, 25, 50, 75, and 100 nM concentrations. Although the percentage of alive cells dropped slightly by increasing the concentration of fingolimod, this decrease did not pass the significance level. Also, there was no variation in obtained result in the time course analyses (Fig. 1, g). Since all of the treated concentrations are below the IC50, based on the previous works, 20 nM concentration was selected for further analysis [29].

### 3.2. *Hotair* and *Malat1* lncRNAs downregulation decreased the *Bdnf* expression

To explore the regulatory role of *Hotair* and *Malat1* lncRNAs on *Bdnf* mRNA expression regulation, their specific DNAzymes were designed to decline their expression. *Hotair* and *Malat1* specific DNAzymes were able to decreased their target mRNAs expression level significantly, Fig. 2a & b (0.18-fold, p= 0.004, and 0.39-fold, p= 0.025, respectively). Then, the *Bdnf* expression level was measured independently and concurrent with *Hotair* and *Malat1* downregulation. The results demonstrated that the expression of *Bdnf* level is affected by *Hotair* (0.19-fold, p= 0.010) and *Malat1* (0.47-fold, p= 0.036) RNA level downregulation (Fig. 2, c).

**Fig. 2.**
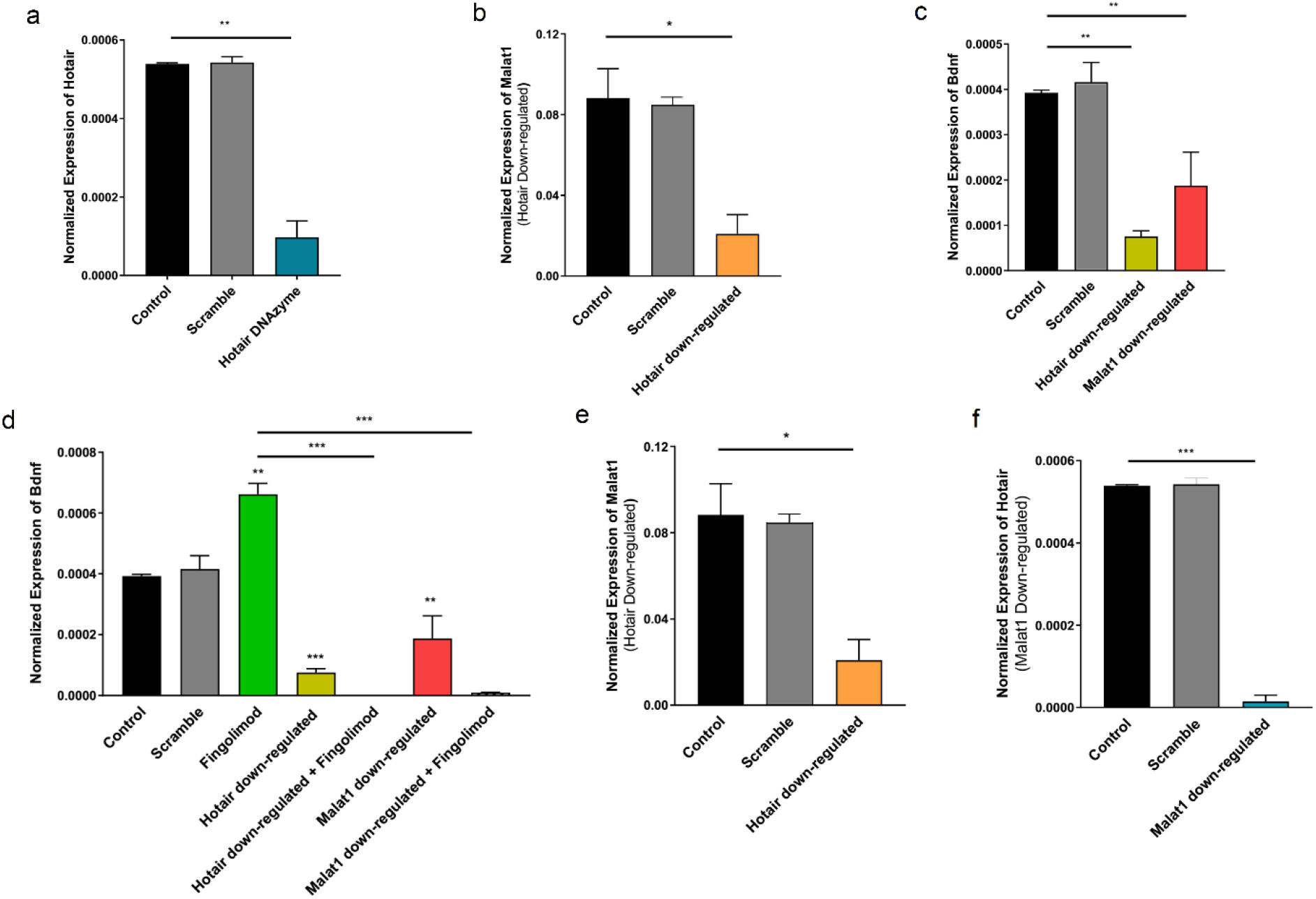
Hotair and Malat1 lncRNAs can regulate Bdnf gene expression. a and Bb, The expression of Hotair and Malat1 lncRNAs was downregulated using their specific DNAzymes. c, Bdnf expression level was affected with reducing Hotair and Malat1 expression level. Obtained result showed that the Bdnf expression reduced due to a decline in expression of these two lncRNAs (unpaired t-test). d, Treatment of mixed glia cells with fingolimod increased the Bdnf gene expression; however, simultaneous downregulation of Hotair or Malat1 with fingolimod treatment not only did not induce BDNF expression but also significantly decreased its expression level. The result suggests that there is a crosstalk between Hotair and Malat1 lncRNAs in regulating Bdnf gene expression. e, a decrease in Malat1 level using its specific DNAzymes reduced the Hotair lncRNA expression level; similarly, f, reducing the Hotair level using its specific DNAzymes lessened the Malat1 expression level (*p< 0.05, **p < 0.01, ***p < 0.001, ****p < 0.0001, unpaired t-test, error bar= standard deviation, n = 3).

### 3.3. *Hotair* and *Malat1* lncRNA downregulation abort the *Bdnf* upregulation by fingolimod treatment

Glial primary culture treatment by fingolimod upregulated *Bdnf* expression (1.7-fold, p= 0.001); however, concurrent fingolimod treatment with downregulation of *Hotair* (0.0015-fold, p<0.0001) or Malat1 (0.013-fold, p<0.0001) lncRNAs not only did not induce *Bdnf* expression but also reduce its mRNA level (Fig. 2, d). This result indicates that *Hotair* and *Malat1* lncRNA play a pivotal role in inducing *Bdnf* through fingolimod treatment.

### 3.4. *Hotair* and *Malat1* lncRNA may regulate each other in a feedback loop

To see if there is any regulatory feedback loop between *Hotair* and *Malat1* lncRNAs, the RNA level of each lncRNA was measured when the other one had been downregulated. The results suggested that a decline in the level of *Malat1* led to downregulation of *Hotair* lncRNA (0.02-fold, p= 0.0005) (Fig. 2, e). Likewise, downregulation of *Hotair* would lead to a decrease in *Malat1* RNA level (0.23-fold, p= 0.03) (Fig. 2, f). These data suggest that there might be a regulatory feedback loop between these two lncRNAs.

### 3.5. OPCs were isolated by the shaking method

The shaking method yielded a highly efficient isolation of OPCs from primary glial culture for further maturation analyses. The whole OPCs purification procedure is depicted in Fig. 3, a. Invert light microscopic images demonstrated the bipolar morphology of isolated OPCs (Fig. 3, b). Immunocytostaining results revealed that the majority of purified culture is consists of Olig2^+^ cells (Fig. 3, c). Quantified immunostaining results represented that the yield of isolation was more than 94% (Fig. 3, d). The purification procedure was repeated at least three times.

**Fig. 3.**
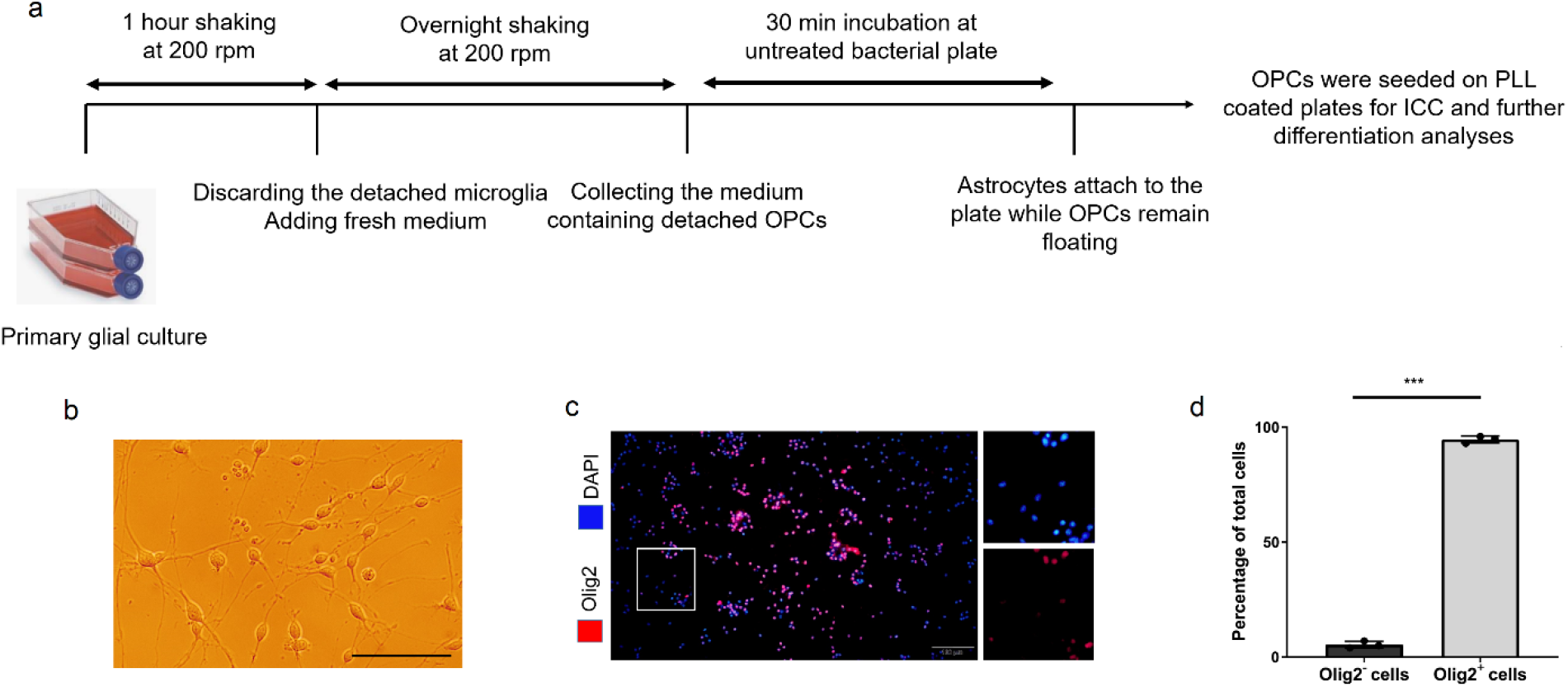
Isolation of OPCs from primary glial culture. a, Procedure for OPCs purification by the Shaking method. b, Light microscopy image represents isolated OPCs with bipolar morphology (scale bar= 100 μm). c, Fluorescence microscopy image demonstrates immunostaining for Olig2^+^ cells (scale bar= 200 μm). d, The graph shows an efficient purification of Olig2^+^ cells from primary glial culture for further analyses (***p< 0.001, paired t-test, n=3). Abbreviation: OPCs= Oligodendrocyte precursor cells, PLL= Poly L-lysin, ICF= Immunocytochemistry.

### 3.6. Fingolimod induced OPCs maturation but not their proliferation

The ICC analysis demonstrated an increasing in MBP^+^ cell after 1 week of fingolimod treatment (Fig. 4, a) compare to the control group (Fig. 4, b). Quantified immunofluorescent signals indicated 2.8-fold increase in the percentage of myelinated oligodendrocytes, which was promoted by fingolimod intervention (p= 0.002, Fig. 4, c). In contrast, the proliferation test did not represent any significant alteration not only in oligodendrocytes proliferation capability but also in their survival in 1 week of fingolimod supplementation (Fig. 4, d).

**Fig. 4.**
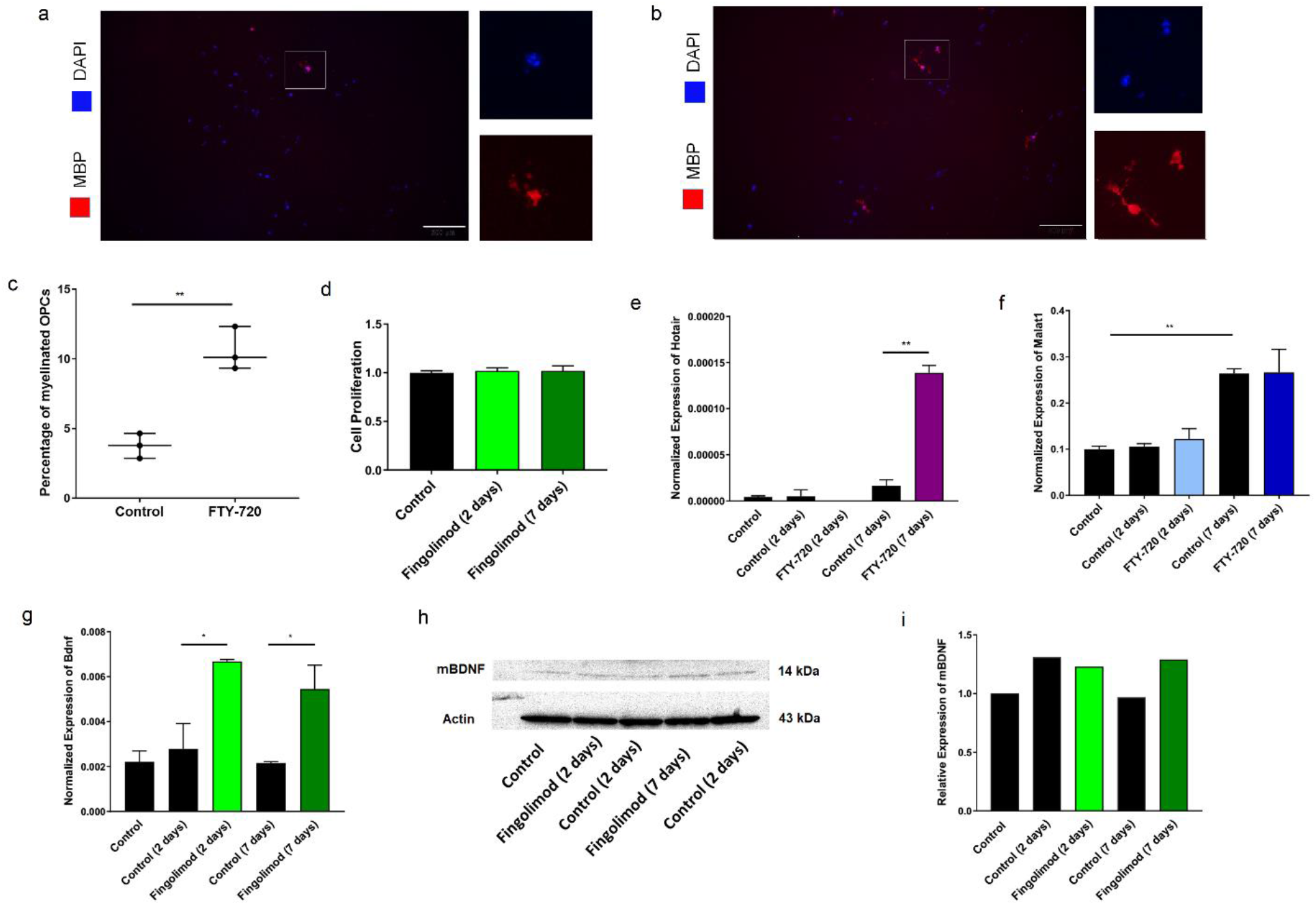
Fingolimod treatment induced OPCs differentiation. MBP/DAPI immunostaining in a, untreated, and b, Fingolimod treated OPCs represented the population of mature MBP^+^ oligodendrocytes (scale bar= 200 μm). c, The quantification of immunostaining results showed that the percentage of Mbp^+^ cells was 3.7% in the control group, while it increased to 10.5% in fingolimod treated OPCs after 1 week (unpaired t-test, error bar= standard deviation, n=3). d, Cell proliferation assay using Resazurin salt indicated that there was no difference in the number of cells between the treated and untreated groups (one-way ANOVA with Tukey’s multiple comparison test, error bar= standard deviation, n=3). e, Fingolimod treatment induced the Hotair expression level after 1 week; however, f, the fingolimod treatment did not alter the Malat1 expression level, but its expression how after 1 week in a differentiation medium ((*p< 0.05, **p < 0.01, unpaired t-test, error bar= standard deviation, n=3). g, The Bdnf gene expression was elevated by fingolimod treatment; while, h, elevation of Bdnf protein expression level by western blot after 2 and 7 days of fingolimod treatment represented; i, Quantitative results of western blot did not show any significant change among different studied time periods. (*p< 0.05, unpaired t-test, n=3). Abbreviation: OPCs= Oligodendrocyte precursor cells.

### 3.7. Fingolimod induced *Bdnf* and Hotair expression along with oligodendrocytes maturation

The fingolimod mechanism of action in inducing OPCs differentiation was analyzed by exploring its possible role in regulating *Hotair*, and *Malat1* lncRNAs and *Bdnf* gene expression. *Hotair* expression upregulated during oligodendrocyte differentiation in response to fingolimod treatment (8.4-fold, p= 0.003, Fig. 4, e); however, its expression was not affected in untreated cells. In contrast to *Hotair*, although *Malat1* expression was not affected by fingolimod treatment, this lncRNA is unregulated along with time in both control and treated cultures (2.5 fold, p= 0.002, Fig. 4, f).

BDNF mRNA and protein expression level were evaluated in control and treated OPCs after 2 and 7 days of fingolimod intervention. RT-qPCR results represented that fingolimod increased *Bdnf* mRNA level significantly after two (2.4-fold, p= 0.03) and seven (2.5-fold, p= 0.04) days of treatment. Without fingolimod treatment, *Bdnf* expression was not raised along with oligodendrocytes maturation at mRNA level. (Fig. 4, g). While, western blot data represented that the expression of mature Bdnf was not altered following fingolimod treatment (P> 0.05, Fig. 4, h & i.

## 4. Discussion

In demyelinating disorders, one of the main problems is that OPCs remain undifferentiated in CNS plaques and do not respond to differentiation stimulators [9]. As a possible therapeutic intervention, fingolimod and Bdnf treatment have shown considerable effects on the plaque remyelination process [1, 2, 5, 6]. Deciphering regulatory pathways involved in OPCs maturation can lead to a better comprehension of demyelinating disorders pathogenesis and facilitates developing new treatments for patients. Moreover, conserved sequences among species is as an indicative for critical preserved functions. Given the conserved sequences of *HOTAIR* and *MALAT1* lncRNAs among humans, mice, and rats [30], these genes may orchestrate conserved and essential regulatory pathways amongst these species.

Previously, it was demonstrated that in cultured neurons, fingolimod treatment induced *Bdnf* expression and hindered neuronal death [31]. Also in EAE mice fingolimod ameliorate memory impairment [32]. Furthermore, in mice Multiple Sclerosis and Alzheimer’ disease models, fingolimod oral administration restored the expression level of *Bdnf* in the cerebral cortices and hippocampi [33]. Similarly, in this study, fingolimod treatment increased *Bdnf* expression in primary glial and OPCs cells.

The obtained results illustrated that fingolimod treatment induces *Bdnf* expression in glial cells, especially astrocytes and microglia. Previous studies also have shown a modulatory role for fingolimod on astrocyte-secreting factors through inhibiting anti-inflammatory cytokines production and stimulating the secretion of BDNF neurotrophic factor, which prevents neural loss [33, 34]. Moreover, fingolimod modulates microglia-secretory factors and switches microglia to M2 phenotype by activating STAT3 pathway and increasing the M2 profile expression, including GDNF, BDNF, IL-10, and TGF-β [35]. Microglia play an essential physiological role in synaptic plasticity, memory, and learning through the production of BDNF [36] indicating their possible role as a double-edged sword in brain. Therefore, fingolimod treatment induces astrocytes and microglial anti-inflammatory and protective roles to maintain the physiologic milieu of the CNS and protect neurons.

Overproduced Bdnf by astrocytes and microglia also play roles in OPCs maturation and myelin repair. In previous studies it was confirmed that Bdnf treatment increases oligodendrocyte proliferation, differentiation, remyelination, neural survival, and axonogenesis [5, 6, 37] through activation of the TrkB receptor downstream signaling pathway [37, 38]. Altogether, fingolimod induces protective aspects of microglia and astrocytes, including anti-inflammatory and neurotrophic factors expression, which in turn have modulatory functions on neural survival and myelin repair.

Given the critical roles of Bdnf in the CNS maintenance, understanding its regulatory systems can be beneficial for finding new therapeutic approaches. Among gene regulatory mechanisms, lncRNAs are an emerging area in RNA research. It is well understood that lncRNAs control different aspects of RNA expression such as, transcription initiation, RNA stability, and translation efficacy in the cytoplasm [16]. In this study, analyzing the *Bdnf* regulatory system demonstrated that a decrease in the expression level of *Hotair* and *Malat1* lncRNAs resulted in lower *Bdnf* expression. This data suggests these two lncRNAs may participate in the fingolimod downstream signaling pathways leading to regulation of target genes, including *Bdnf* through a lncRNA-mediated mechanism such as the competitive endogenous RNA (ceRNA). The competitive endogenous RNA (ceRNA) hypothesis can interpret this association among Bdnf expression and *Hotair* and *Malat1* lncRNAs. The ceRNA hypothesis represents the interaction among miRNAs, lncRNAs, and mRNAs through microRNA binding sites (MREs) [39].

Simultaneous *Hotair* and *Malat1* lncRNAs downregulation under fingolimod treatment, not only did not enhanced *Bdnf* expression but also decreased its expression level compare to DNAzyme treated groups through an unknown mechanism.

Also, this study showed that there might be a regulatory feedback loop between *Hotair* and *Malat1* lncRNAs. Downregulation of each lncRNA results in reducing the expression level of the other one. The ceRNA hypothesis can interpret this positive regulatory feedback loop between *Hotair* and *Malat1* lncRNAs. This hypothesis indicates that lncRNAs and mRNAs regulate each other by competing to scavenge for the restricted miRNA pool. Hotair may regulate *Malat1* by sponging miR-217, which has a confirmed binding site on *Malat1* sequencing [40]. Furthermore, the downregulation of *Malat1* enhances mature miR-1 expression even more than its precursors. The existence of the miR-1 binding site on *Malat1* lncRNA indicates that this lncRNA increases *Hotair* level through sponging miR-1 [41]. CeRNA hypothesis may justify the observed positive regulatory feedback loop between these two lncRNAs.

*Malat1* is expressed ubiquitously and its expression level is higher than many coding genes, even comparable to a highly transcribed β-Actin housekeeping gene [30]. On the contrary, *Hotair* is a very low abundant lncRNA not only in normal tissues but also in different diseases [42]. These findings are in line with our results. The issue to consider is that a lncRNA abundance should not be misunderstood with its importance. As for the low expressed *Hotair*, there is accumulating evidence for its prominent roles in development, chromatin modification, pathogenesis, and creating RNA scaffold, to mention some [43].

More than stimulatory effects on *Bdnf* expression, fingolimod also can increase myelin formation [14, 15, 29, 32]. The obtained data showed that fingolimod increase OPCs differentiation, however, the MBP^+^ cells are present in the untreated group with fewer number. After 7 days, 3.7 percent of OPCs in the differentiation medium expressed MBP marker without fingolimod treatment. In another study, after 12 days of OPCs maintenance in differentiation medium, the expression level of PDGFαR reduces by half and the MBP was expressed in about 20 percent of total cells [44]. OPCs can differentiate in differentiation medium without any further stimuli; however, different substances can facilitate this process and make it more efficient. Here, it was showed that adding fingolimod to the differentiation medium have stimulatory effect on MBP expression. As represented by previous studies, Triiodo-L-thyronine (T3), which is one of the differentiation medium ingredients, controls the specification and differentiation of OPCs to the mature oligodendrocytes. T3 promotes and helps the timing of precursor cell differentiation through a currently unknown mechanism [44, 45].

Here, it was demonstrated that fingolimod induces OPCs maturation, but not their proliferation. Given the fact that differentiation and proliferation are two contradictory mechanisms [46], this data indicates that fingolimod hinders OPCs proliferation to trigger the differentiative pathways. Fingolimod treatment stimulated OPCs differentiation and MBP expression concomitant with increasing *Bdnf* mRNA. Elevated *Bdnf* level subsequent to fingolimod treatment has demonstrated in previous researches [31, 47, 48]. Also, protective role of Bdnf in the CNS has been clearly declared [1, 2, 5, 6]. Altogether, OPCs undergo maturation by fingolimod treatment, and the Bdnf neurotrophic factor may play role in their differentiation process. Unchanged mature Bdnf protein expression during OPCs differentiation was observed may be due to the pro-Bdnf process in the secretory vesicles, so the elevation of mature-Bdnf cannot be detected because the mature Bdnf is secreted immediately from the cell [47].

More than *Bdnf*, seven days of fingolimod treatment enhanced the expression of *Hotair* simultaneously with MBP expression. Hence, this result suggests that long-term of fingolimod treatment induces *Hotair* expression in OPCs which may play a role in their differentiation process. Dissimilar to *Hotair* lncRNA, *Malat1* expression increased independently from fingolimod treatment, indicating that *Malat1* lncRNA may play an intrinsic role in the OPCs maturation in differentiation medium. Another interpretation for this hesitated response is the selected time window [49]. There is a possibility for a short-term response of *Malat1* and Hotair in response to fingolimod treatment, so they cannot be assessed readily.

Upon fingolimod treatment, increased *Bdnf* expression did not match with alteration in *Malat1* and *Hotair* lncRNAs expression level. Subsequently, unlike glial culture, there is no correlation between *Bdnf* and lncRNAs expression levels. Given that microglia and astrocytes constitute the majority of the primary culture, this observed correlation in the downregulation experiment belongs to astrocytes and microglia cells. Future studies on animal models of demyelinating disorders could provide valuable data on the *Malat1* and *Hotair* lncRNAs mechanism of action in the remyelination process.

## 5. Conclusion

The results of the current study showed that *Bdnf* regulatory mechanism probably is distinct in different types of glial cells. As mentioned above, *Hotair* and *Malat1* may control *Bdnf* expression level significantly, in primary glial cells. Also, it was demonstrated that *Hotair* and *Bdnf* may play role in fingolimod-induced OPCs differentiation. Nevertheless, *Malat1* takes part as a natural modulator in OPCs differentiation. Given the obtained results, *Hotair* and *Malat1* may play critical regulatory role in fingolimod mechanism of action, *Bdnf* expression and oligodendrocytes differentiation.

## Declarations

## Acknowledgment

The authors thank Javad Mirnajafi-Zadeh for providing rat pups for this study. We also thank the funding source for their support.

## Author information

### Contributions

All authors contributed to the study conception and design. Material preparation, data collection and analysis were performed by Fatemeh Khani-Habibabadi, Mehrdad Behmanesh, Leila Zare, and Mohammad Javan. The first draft of the manuscript was written by Fatemeh Khani-Habibabadi and all authors commented on previous versions of the manuscript. All authors read and approved the final manuscript.

### Corresponding authors

Correspondence to M. Behmanesh and M. Javan.

## Ethics declarations

### Ethics approval

The procedures of this research were approved by the ethics community of Tarbiat Modares University (ID: IR.TMU.REC.1396.607).

### Availability of data and material

The data that support the findings of this study are available on request from the corresponding author.

### Code availability

Not applicable.

### Consent to participate

Not applicable.

### Consent for publication

Not applicable.

